# Monoacylglycerol disrupts Golgi structure and perilipin 2 association with lipid droplets

**DOI:** 10.1101/2021.07.09.451829

**Authors:** Lydia-Ann L.S. Harris, James R. Skinner, Trevor M. Shew, Nada A. Abumrad, Nathan E. Wolins

**Affiliations:** bioMérieux Inc., 595 Anglum Rd, St. Louis MO 63042; Department of Medicine, Center for Human Nutrition, Washington University School of Medicine, St. Louis, Missouri 63110; Cell Biology and Physiology, Washington University School of Medicine, St. Louis, Missouri 63110

**Keywords:** Lipid absorption, enterocyte, coatomer, fractionation, OP9, Brefeldin A

## Abstract

The two major products of intestinal triacylglycerol digestion and lipoprotein lipolysis are monoacylglycerols (MAG) and fatty acids. In the gut, these products are taken up by enterocytes and packaged into perilipin-coated cytosolic lipid droplets and then secreted as chylomicrons. We observed that fat feeding or intragastric administration of triacylglycerol oil caused the enterocyte Golgi to fragment into submicron puncta dispersed throughout the cytosol. Further, this apparent Golgi dispersion was also observed in cultured fibroblasts after treatment with fat (cream) and pancreatic lipase, but not when treated with deactivated lipase. We therefore hypothesized that a hydrolytic fat product, specifically monoacylglycerols, fatty acids or a combination of these molecules can trigger Golgi fragmentation. Disruption of coatomer function is known to cause Golgi to fuse with the ER, and blocks perilipin 2 delivery to lipid droplets. Thus, we assessed the effects of MAG on coatomer distribution, Golgi structure and perilipin 2 localization. We found that MAG, but not fatty acids, dispersed coatomer from the Golgi, fragmented the Golgi and caused perilipin 2 to accumulate on cellular membranes. Thus, our findings suggest that monoacylglycerol production during digestion disperses the Golgi, possibly by altering coatomer function, which may regulate metabolite transport between the ER and Golgi.

## INTRODUCTION

Chronic inactivity and overeating increase fat stores and hasten the onset of metabolic disease. Interestingly, short-term overeating can mute metabolic signals and drive ectopic lipid accumulation. Conversely, caloric restriction even before significant weight loss can reverse overeating-induced metabolic derangement [1–3]. These observations suggest that lipids being processed, rather than stored fat, determine energy metabolism. Dietary fat-derived molecules transmit signals, fuel cells, function in vesicular transport and are constituents of the membranes that surround cells, organelles and lipid droplets (LDs) [4]. These roles raise the possibility that high levels of dietary fat- derived molecules might alter the signaling and transport processes that direct them to fat droplets.

The coatomer refers to the protein complexes COPI and COPII. These complexes are involved in the assembly of transport vesicles that move between the ER and Golgi. COPI facilitates transport from the Golgi to endoplasmic reticulum, and COPII from ER to the Golgi. The coatomer and other trafficking coat complexes regulate vesicle formation, and shape intracellular and plasma membranes [5–7]. Predictably, modulating trafficking coats quickly and profoundly perturbs the cellular membrane and consequently cellular architecture. [5, 8, 9]. Since TAG and its hydrolytic products partition into and thus travel with membranes, metabolism and membrane trafficking are linked. For example, inhibiting membrane traffic with the coatomer-disrupter brefeldin A (BFA) fuses the Golgi and endoplasmic reticulum (ER) into one compartment and inhibits trafficking of perilipin 2 and ATGL to LDs [10, 11, 12]. The ER membrane is where most fatty acid esterification occurs and the point at which fat is directed either to cytosolic LDs or to distal secretory compartments for lipoprotein assembly and secretion [13]. Coatomer controls membrane traffic between the ER and Golgi [14]. In addition to controling vesicle traffic from the Golgi back to the ER, COPI controls protein traffic from ER to LD, and buds tiny LDs from larger artificial triacylglycerol droplets [8, 12, 15, 16]. The proteins COPI traffics to lipid droplets include perilipin 2 and adipocyte triglyceride lipase [12, 17, 18]. Interestingly, COPI does not traffic perilipin 3 to LDs [12]. By directing trafficking between ER, LD and distal secretory compartments, coatomer plays a role in cellular fat processing.

In the intestinal lumen, ingested fat is emulsified into particles, and triacylglycerols (TAGs) in these particles are hydrolyzed by pancreatic lipase and the products—fatty acid and 2-monoacylglycerol (MAG)—taken up by enterocytes [19, 20]. Uptake of 2-MAG by enterocytes has been shown to influence feeding by controlling enterocyte hormone release [21]. Further, high MAG levels, due to pharmacological or genetic factors, affect behavior and metabolism. Specifically, endocannabinoids are MAGs [22, 23]. MAG is amphipathic and thus will partition into membranes. However, uptake of 2-MAG and its potential effects on lipid signaling, membrane trafficking and processing during lipid absorption are unknown [24]. In this study we examined if high MAG influences Golgi structure and that of the LD membrane. We find that after a TAG gavage, the Golgi in enterocytes is dispersed. Golgi integrity and perilipin 2 targeting to LD are dependent on COPI components of coatomer, and we show that in cultured cells they are disrupted by 2-MAG and that 2-MAG disperses the COPI coat. Our data suggest that 2-MAG regulates protein and membrane trafficking by modulating formation of the COPI complex.

## MATERIALS AND METHODS

### Reagents

2-oleoylglycerol was purchased from Avanti Polar Lipids, Sigma-Aldrich (St. Louis, MO) and Santa Cruz Biotechnology. Bovine serum albumin was from Equitech-Bio (Kerrville, TX, catalog number BAH66). Other regents were from Sigma-Aldrich unless stated otherwise. Oleic acid was complexed to albumin as described previously [25].

### Antibodies

Monoclonal antibody against mouse Golgi matrix protein GM130 was purchased from BD Biosciences; rabbit polyclonal antibody against Golgi SNARE protein (GS28 Antibody) from Proteintech™ (Chicago, IL, catalog number 16106-1-AP); rabbit polyclonal against β-COP from Thermo Scientific (catalog number PA1-061) and rabbit polyclonal antibody against calnexin from Enzo Life Sciences, Inc. (Farmingdale, NY, catalog number ADI-SPA-860). Guinea pig antiserum against perilipin 2 was purchased from Fitzgerald Industries (Concord, MA, catalog number RDIPROGP40) and rabbit polyclonal antibody against perilipin 3 was raised using the immunizing peptide CSGPFAPGITEKTPEGK. Rabbit polyclonal antibody against perilipin 2 was previously described [26].

### Immunohistochemistry (IHC)

The small intestine was harvested from overnight fasted C57BL/6 mice or from mice 2 h after refeeding high fat food. The mice were euthanized with carbon dioxide. All protocols were approved by the Animal Care Committee of Washington University. The small intestine 3 cm proximal to the stomach was excised and cut longitudinally, opened, pinned flat and fixed. Intestines were embedded in agarose before paraffin embedding to position tissue so that enterocytes are cut longitudinally. Sections were stained and imaged as described previously [27].

### Transmission Electron Microscopy

Intestines were harvested and pinned flat as described for IHC and then paraformaldehyde/glutaraldehyde fixed. Fixed intestines were processed with osmium tetroxide and uranyl acetate followed by resin embedding and sectioning. Intestine samples were imaged at the Molecular Microbiology Imaging Facility at Washington Univ. in St. Louis on a JEOL 1200 transmission electron microscope (JEOL USA Inc.) [27].

### Cell culture

C2C12 and COS7 cells were cultured in Dulbecco’s modified eagle medium (DMEM) with 10% fetal bovine serum supplemented with antibiotics. OP9 mouse stromal cells were cultured as described previously [28].

### Cream hydrolysis

Heavy cream (~ 40% fat by weight) was treated for 10 min while shaking at 37°C with 10 mg/ml native or boiled (10 minutes) lipase from porcine pancreas from Sigma (St. Louis, MO; catalog number L3126).

### Immunofluorescence microscopy (IF)

Cells were fixed, stained and imaged as described in Harris et al. [29].

### Treatments with lipids and BFA

Cells were incubated at 37°C rotating at 60 rpm in DMEM with 10% fetal bovine serum with antibiotics and buffered with 50 mM HEPES. MAGs and BFA were delivered from stocks dissolved in ethanol, 20 mg/ml and 5 mg/ml, respectively.

### Cell counting

OP9 cells were treated and co-stained with GM130 and β-COP as described in Figure 3. Fluorescent signal was quantified using ImageJ. The fraction of signal in the Golgi region was calculated by dividing fluorescent signal in the Golgi regions by the total fluorescent signal.

### OP9 cell fractionation

OP9 cells were grown in 0.4 mM albumin bound oleate to induce perilipin 2 protein. These cells were fractionated into pelleting, soluble and floating fractions, as described previously [30].

### Thin layer chromatography

Fractions (200 μl) were extracted and the lipids dissolved in hexane:ether (1:1), loaded on Whatman Partisil LK6D Silica Gel TLC plates and resolved in hexane:ether:acetic acid (70:30:1.2) then stained with molecular iodine as described previously [25, 31].

## RESULTS

### The Golgi in enterocytes disperses during fat absorption

To maximize dietary fat absorption, enterocytes transiently store fat in LD and transport it into the lymph as chylomicrons [32]. Due to their hydrophobicity, molecules hydrolyzed from fat partition into or between membrane leaflets. It is unknown how dietary fat absorption alters enterocyte membrane and fat droplet architecture to facilitate intracellular lipid trafficking and storage. To this end, we tested the effect of fat ingestion on the membrane structures of Golgi and LD in duodenal enterocytes isolated from mice after either an overnight fast or 2-hours after a 10 μl/g corn oil gavage (Figure 1). In fasted mice enterocytes had well defined Golgi, evidenced by intense, juxtanuclear GM130 (Golgi) staining (arrows). Perilipin 3 staining was diffuse indicating a lack of nascent lipid droplets [33–35]. In contrast, the enterocytes of fat-fed mice, which featured large perilipin 3-stained lipid droplets, lacked the ordered GM130 staining (Figure 1A). Ultrastructural studies showed similar results. Similarly, the intact Golgi structure was evident in the duodenum from fasted mice, but LDs were not, while in fat-gavaged mice the Golgi appeared diffuse as much of the cytosol was fat filled (Figure 1B). These observations show that during fat absorption the Golgi in enterocytes disperses, as perilipin 3 coated LDs emerge.

**Figure 1.**
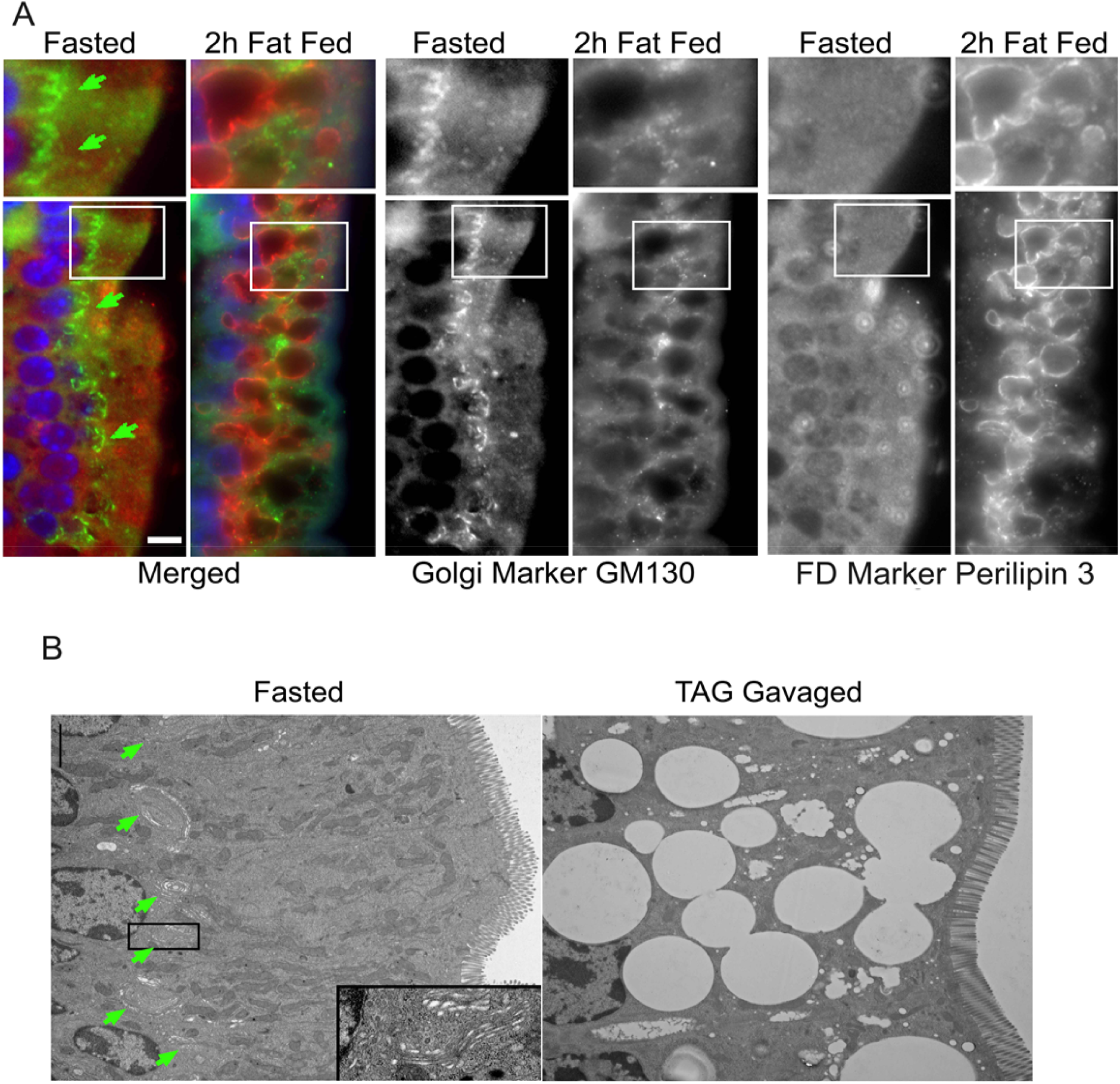
High fat feeding causes Golgi dispersion in mouse enterocytes. Panel A. Shows intestine from mice overnight-fasted (Fasted) or overnight-fasted then re-fed high fat food for 2h (2h Fat Fed). Intestines were stained for a Golgi marker, GM130, and a marker for nascent fat droplets (FD), perilipin 3. For more details see MATERIALS AND METHODS. Anti-GM130 was used at 1.25 μg/ml (shown as green) and perilipin 3 antiserum was diluted 1:30 (shown as red). Green arrows point to Golgi. Bar = 10 μm. Panel B. Electron micrographs of mouse enterocytes. Mice were overnight-fasted. Intestines were either harvested without further treatment (Fasted) or 2h after a corn oil (10 μl/g of mouse) gavage (TAG Gavaged). Intestines were prepared as described in the MATERIALS AND METHODS. Green arrows point to Golgi. Bar = 5 μm (upper left).

Dietary fat absorption is regulated by molecules released from chyme, and by hormonal and systemic signals [36]. We hypothesized that uptake and storage of molecules released from chyme cause the changes we observed in cellular architecture. To exclude the effects of systemic signals, we treated OP9 fibroblast cells with pancreatic lipase-treated milk fat. We found that milk fat treated with active lipase, but not heat-killed lipase, causes Golgi dispersal in OP9 fibroblasts (Figure 2) and COS7 cells (not shown). These observations indicate that a hydrolytic product or products of milk fat cause the Golgi to disperse. The dispersed Golgi formed a cytosolic reticulum that has significant overlap with calnexin-stained membrane, indicating Golgi fusion with the ER (Figure 2).

**Figure 2.**
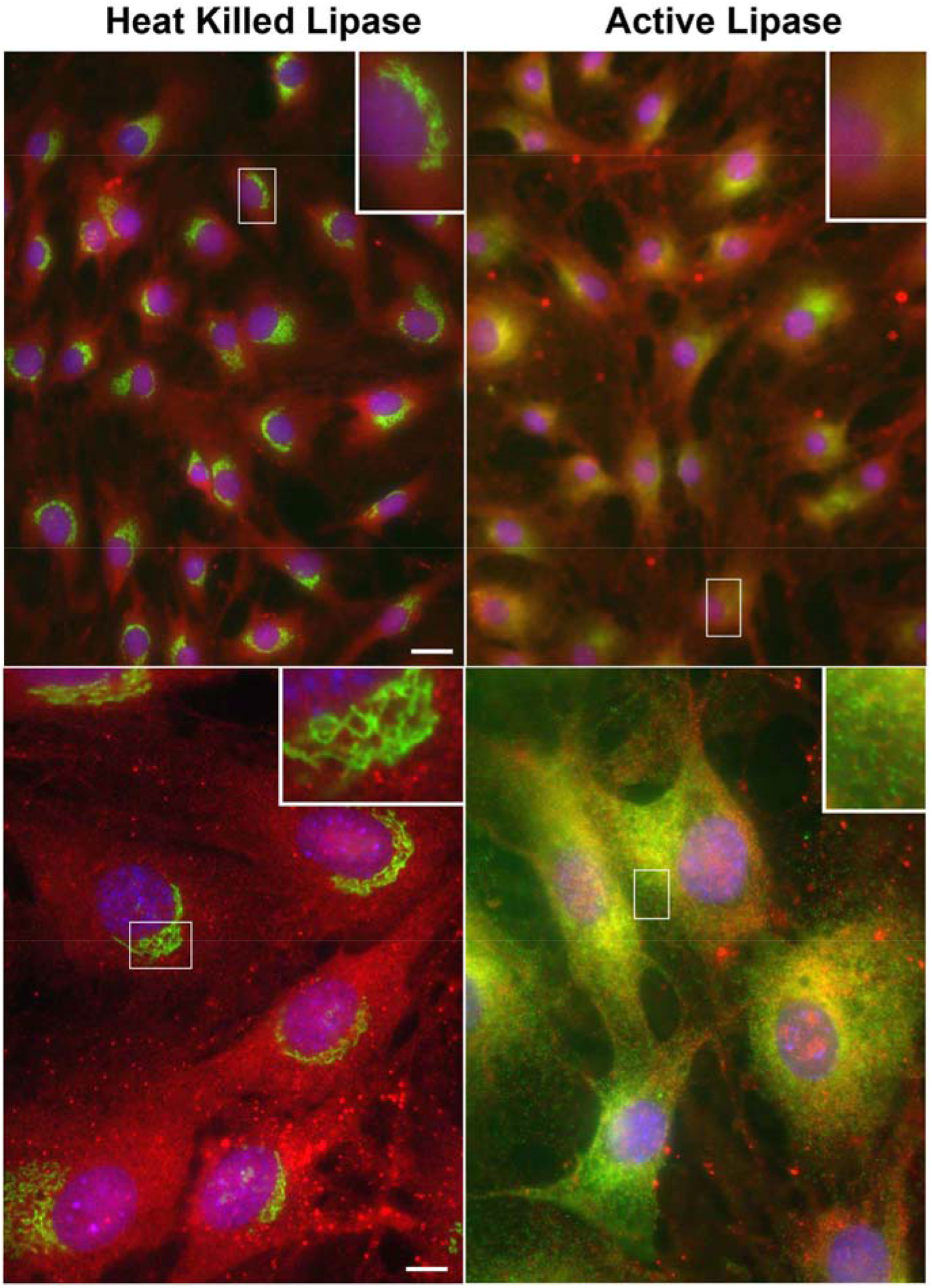
Hydrolyzed milk fat causes Golgi and ER to fuse. OP9 fibroblasts were incubated with heavy cream for 10 min. The heavy cream was treated with boiled or native pancreatic lipase diluted 10-fold in culture media. Fibroblasts were fixed and stained with GM130 at 500 ng/ml (green) and calnexin 1:500 (red). See MATERIALS AND METHODS for more details. Bar = 30 μm in top panels and 10 μm in bottom panels.

### Long chain MAGs cause Golgi ER fusion (GERF)

Pancreatic enzymes hydrolyze TAG to MAG and FA. Since gavaged corn oil causes the Golgi to disperse (Figure 1B), we infer that MAG, FA or MAG in combination with FA causes GERF. To determine which molecule or molecules causes GERF, we tested MAG 2-oleoyl-glycerol, or albumin bound-oleate, and the two together in OP9 fibroblasts and C2C12 myoblasts (Figures 3–6). The MAG 2-oleoyl-glycerol induced GERF as indicated by the co-localization of signals for Golgi and ER markers, and this effect was also observed in CHO and COS7 cells (not shown). Further, almost all cells were affected, as shown by all treated cells having less signal in the Golgi regions than vehicle control cells. COPI signal was similarly affected (Figure 3B). Using two different antibodies and four cell types we show that MAG induces GERF (Figures 3,4, and 6).

**Figure 3.**
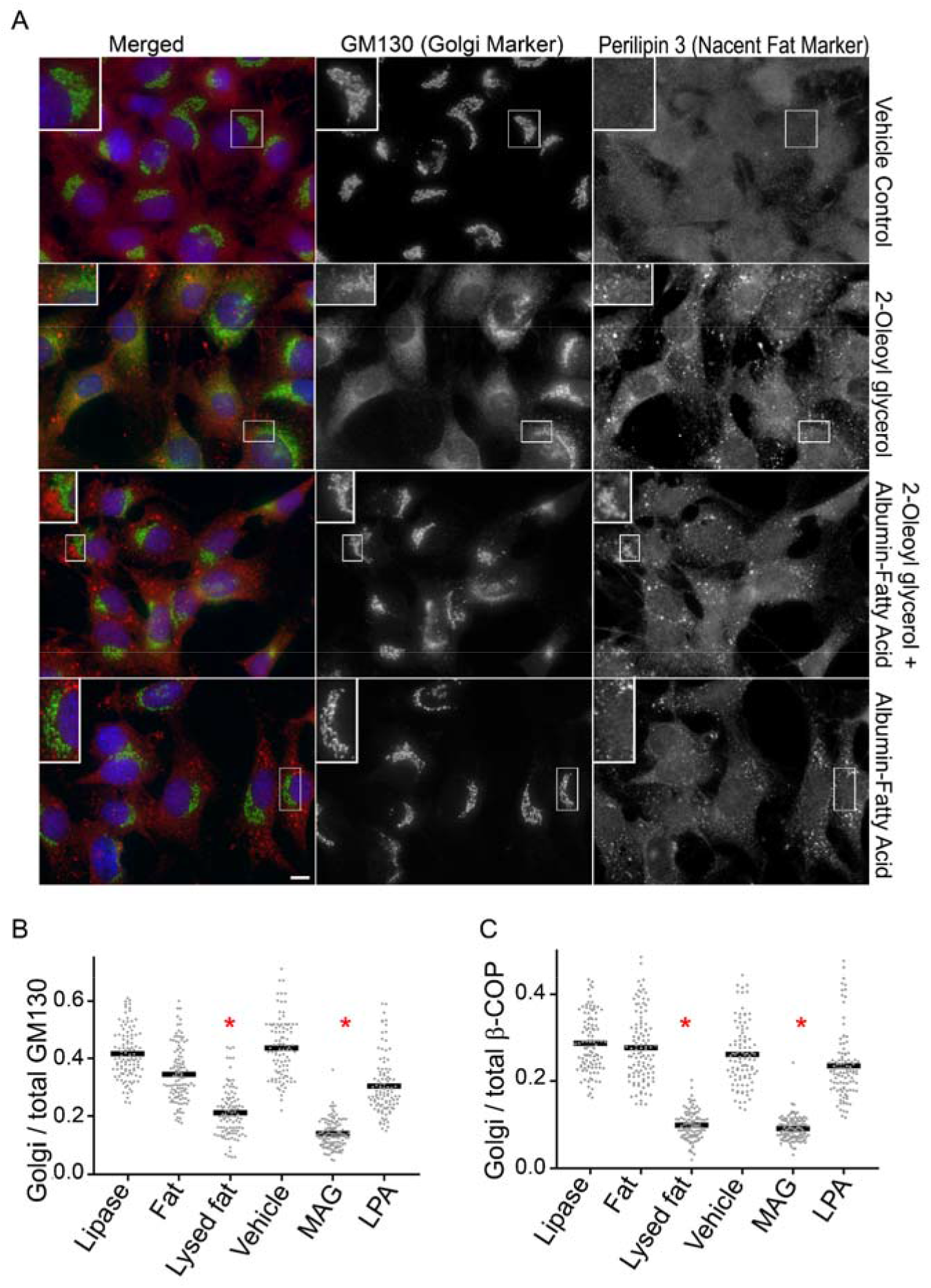
Oleoylglycerol causes Golgi and ER to fuse. (A) Shown are OP9 cells treated with combinations of Vehicle, Albumin-Fatty Acid and MAG, and stained with antibodies against a Golgi marker (GM130 shown as green) and a nascent fat-binding protein (perilipin 3 shown as red). Cells were treated with vehicle, 1.8 mM albumin fatty acid and 60 μg/ml 2-oleoylglycerol for 30 min as indicated. Bar = 10 μm. (B) OP9 cells were treated with 20 μg/ml 2-oleoylglycerol (MAG) for 30 min. Cells were fixed and stained using 1.3 μg/ml anti-GM130 and β-COP (1:400). The antibody signals in the Golgi and the whole cell were quantitated. The fraction of signal in the Golgi was calculated by dividing the signal in the Golgi region by the signal in the whole cell. Data from anti-GM130 and β-COP were analyzed separately. As indicated by *, ANOVA shows that MAG and Lysed fat treatments resulted in significantly less signal in the Golgi than Vehicle for both anti-GM130 and β-COP (P < 0.01). For more details see MATERIALS AND METHODS.

**Figure 4.**
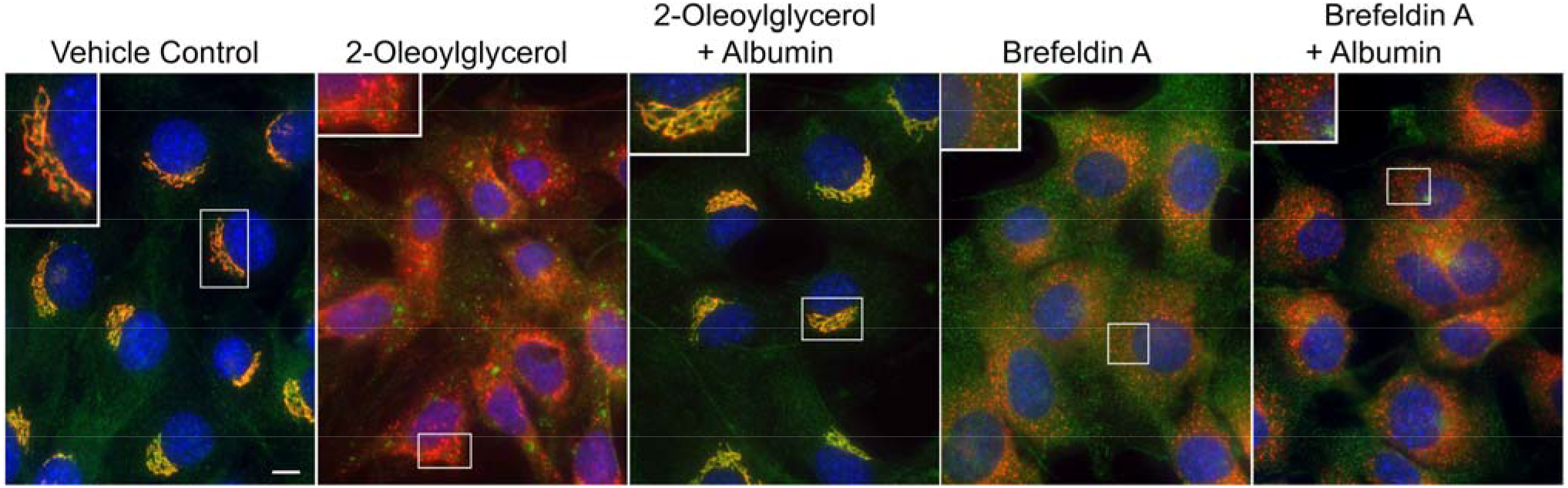
Albumin blocks MAG-induced, but not BFA induced, Golgi and ER fusion. Shown are OP9 cells treated with combinations of MAG, BFA and albumin. Cells were treated for 30 min with 40 μg/ml 2-Oleoylglycerol (MAG), 10 μg/ml BFA, and 2.2 % albumin, as indicated. Cells were stained with the Golgi markers anti-GM130 (shown as green) and anti-GS28 (shown as red) using 750 ng/ml and 2.5 μg/ml, respectively. Bar = 10 μm.

Glycerol and FA are released from fat cells when energy needs are not met by food. In contrast, MAG and FA are released from dietary fat in the intestine by pancreatic lipase and then from lipoproteins by lipoprotein lipase [37–39]. Thus, abundant MAG indicates fat ingestion. We found that MAG induced GERF, whereas FA did not. These observations suggest induction of GERF by MAG plays a role in transporting fat to storage (Figure 3). To further illustrate the role of MAG, but not FA, in GERF, we treated cells with albumin-bound FA, which surprisingly also inhibited GERF (Figure. 3A). However, a review of the literature revealed that albumin binds MAG and that this binding is not displaced by the presence of fatty acids [40]. This led to the hypothesis that albumin blocked MAG-induced GERF by lowering the free MAG concentration. Consistent with this hypothesis, we found that albumin with or without FA blocks MAG induced GERF (Figures 3 and 4, respectively) and FA without albumin does not (not shown). Further, albumin did not block BFA induced GERF (Figure 4).

Several lines of evidence support the notion that BFA induces GERF by causing COPI complex shedding from Golgi membranes [41, 42]. One observation supporting this notion is that BFA disperses the COPI complex within seconds, which is before GERF [41]. We find that MAG, like BFA, rapidly disperses the β-COP subunit of the COPI complex (<30 s) before MAG induced GERF occurs, which is shown by GM130 signal still being largely Golgi associated while β-COP is dispersed (Figure 5). These data suggest that MAG induces GERF by attenuating COPI function.

**Figure 5.**
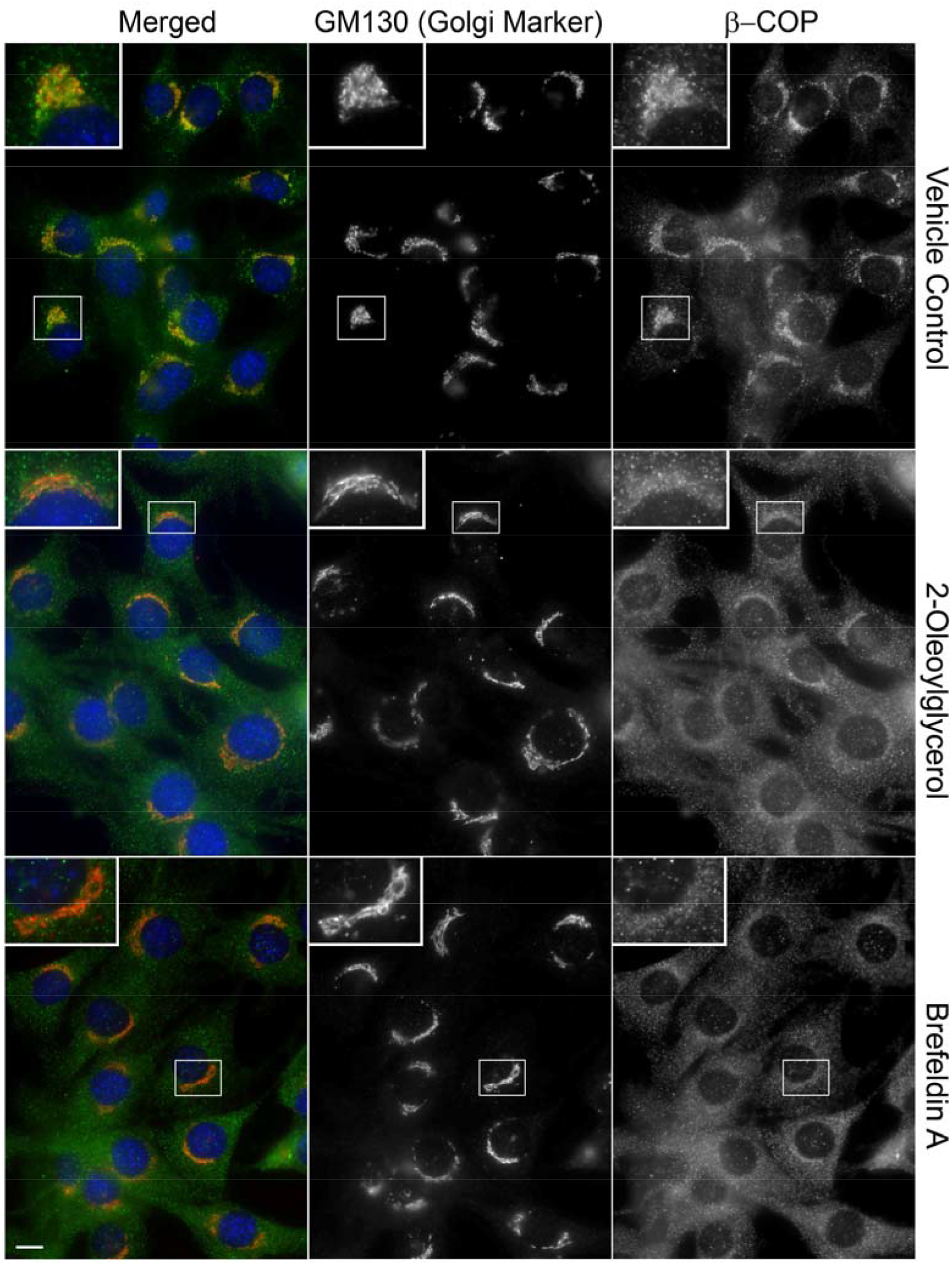
MAG rapidly dislocates β-COP from the Golgi. C2C12 myoblasts were treated with 20 mg/ml 2-oleoylglcerol for 30 s and fixed. Myoblasts were then stained with GM130 (1.3 μg/ml anti-GM130) and β-COP (1:400). Bar = 10 μm.

### MAG modulates the association of perilipin 2 with fat droplets

The coatomer directs protein trafficking between ER and Golgi, and ER and LD. A specific example of this control is that coatomer is required to traffic perilipin 2 to LDs [12]. Thus, coatomer directs fat to either cytosolic droplets for storage or lipoprotein particles for export [12, 43–46]. Release of MAG by hydrolysis of extracellular fat particles after feeding is an integral component of postprandial fat metabolism [37, 38, 47]. We tested the hypothesis that MAG may modulate coatomer function and the delivery of perilipin 2 to LDs based on the report that BFA disruption of the coatomer coat both drives GERF and inhibits perilipin 2 delivery to FDs [12]. To test the effect of MAG treatment on coatomer function, we grew OP9 cells with physiologic levels of fatty acid to induce detectable levels of perilipin 2, and treated these cells with either vehicle or MAG. Next, we fractionated or imaged cells. Immunoblotting of fractions revealed that MAG treatment increased perilipin 2 levels in the pellets (membrane fraction) [Figure 6] and imaging showed that MAG increased perilipin 2 reticular staining, suggesting retention of perilipin 2 in the ER (Figure 6). Thus MAG, like BFA, inhibits perilipin 2 delivery to LDs [12]. Further, MAG does not inhibit perilipin 3 delivery to LD (Figure 1), consistent with the report by Soni et al. [12] that perilipin 2 but not perilipin 3 delivery to FD is coatomer dependent. The observations that MAG within seconds dissociates β-COP from the Golgi and attenuates perilipin 2 LD association suggest that MAG modulation of coatomer function contributes to GERF.

**Figure 6.**
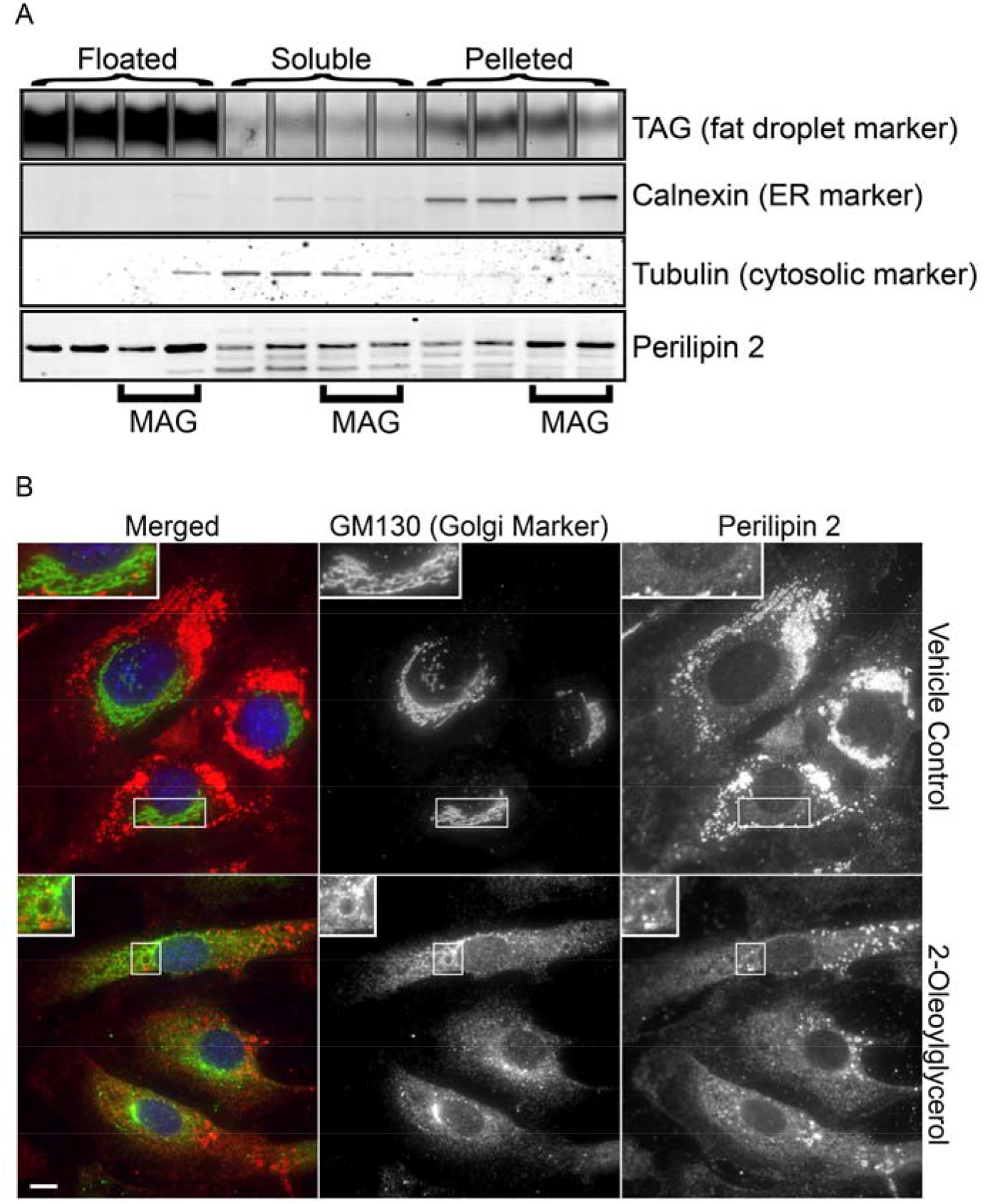
MAG modulates the association of perilipin 2 with fat droplets. (A) Shown are immunoblots of vehicle or MAG-treated OP9 cell fractions. 2-100 mm plates of cells (2 replicates for each condition) were treated with ethanol (vehicle control) or with 60 μg/ml of the MAG 2-Oleoylglycerol, as indicated. After 40 min at 37°C, cells were fractionated into floating, soluble and pelleting fractions. Proteins were extracted from each fraction and equal portions of each extract were immunoblotted and probed with the antibody indicated. Lipids (incl. TAG) were also extracted from equal portions of each fraction, then resolved by TLC and stained with molecular iodine. Anti-calnexin and -tubulin were used at 1 μg/ml and rabbit anti-perilipin 2 at 2 μg/ml. (B) Shown are vehicle or MAG treated OP9 cells stained for perilipin 2 and the Golgi marker GM130. OP9 cells were treated as described in (A) and then fixed. Anti GM130 was used at 500 ng/ml (green) and Guinea pig antiserum to perilipin 2 at 1:1000 (red). Bar = 10 μm.

## Discussion

Digestion of dietary TAG by pancreatic lipase in the intestinal lumen yields FAs and MAG for enterocyte uptake and processing. The FAs are esterified in the ER to regenerate TAG and make other complex lipids [37, 47–49]. From the ER, coatomer mediated trafficking directs the lipids to membrane bilayers, cytosolic droplets or secreted lipoproteins. We show that the elevation of MAG after fat feeding triggers changes in coatomer dependent trafficking. Specifically, MAG induces GERF (Golgi-ER fusion) that is similar to BFA induced GERF in both time course (<30 min.) and morphology (Figures 3,4, and 6). In addition, we show that the coatomer-dependent targeting of perilipin 2 to LD [12] is attenuated by MAG (Figure 6). The observed effects of MAG appear to be limited to the ER, Golgi and LD. Other organelles and structures including lysosomes, nuclei, mitochondria and the actin cytoskeleton do not appear affected (not shown).

MAG is also generated during TAG hydrolysis by lipoprotein lipase in the circulation. Consistent with MAG being released from lipoproteins, serum albumin binds MAG [40] and MAG bioavailability in the circulation is buffered by albumin [40, 50–53]. Unsurprisingly, we observe that high serum albumin concentration abolishes MAG-induced GERF (Figure 4). When lipoproteins dock to endothelial cells, MAG is delivered by apical surface lipoprotein lipase. Presumably, uptake, dilution, and albumin binding limit MAG boluses temporally and spatially. This suggests MAG boluses are delivered to specific sets of cells after fat feeding. This leads us to conjecture that MAG released from dietary fat serves as a signal for subsequent lipid and membrane trafficking.

Numerous lipids can modulate membrane trafficking [54–58]. In particular, elevated levels of lysophosphatic acid (LPA) in the Golgi was shown to cause Golgi-ER fusion [54, 57, 59]. Since MAG can be converted to LPA, this conversion and consequent accumulation of LPA in the Golgi may contribute at least in part to the effects we observed. However, in addition to LPA, various glycerolipids are implicated in membrane trafficking [25, 26, 60–64]. However, the GERF inducing lipid in our experiments likely has a single acyl chain, since the acylation inhibitor triacsin C did not inhibit MAG induced GERF (not shown).

What is the role of MAG-induced Golgi-ER fusion? We speculate that it directs absorbed fat to lipoprotein formation and facilitates the process. Chordates move dietary fat through a series of particles in enterocytes, lymph, blood, liver and blood again before storing the fat in adipocyte droplets. We show that MAG, like coatomer-disrupting BFA, causes GERF in cultured cells and attenuates perilipin 2 LD association [10–12] (Figures 3, 4, and 6). We also find, that like BFA, MAG rapidly dissociates β-COPI from the Golgi preceding the induction of GERF [41] (Figure 5); thus disrupting the coatomer coat, which causes GERF [65–67], likely mediates the effect of MAG. We speculate that it functions to shape coatomer-dependent trafficking to streamline fat uptake and lipoprotein secretion. Consistent with this, treatment with GERF inducing BFA increases fatty-acid labeling of cholesteryl ester in lipoproteins by CaCo2 and HepG2 cells [68]. Also, apoB overexpression causes the ER to wrap around FD and BFA increases this wrapping while it attenuates perilipin 2 association with FDs. In contrast, overexpressing perilipin 2 decreases apoB-ER wrapping [69]. These findings together with our current data suggest that MAG inhibition of coatomer and fat traffic to LD directs the fat retained in the ER to lipoprotein generation.

## Abbreviations used

TAG: triacylglycerol
MAG: monoacylglycerol
ER: endoplasmic reticulum
GERF: Golgi-ER fusion
BFA: Brefeldin A

## Acknowledgments

The authors acknowledge the technical help of Terri Pietka. We also thank Drs. Daniel S. Ory, Kimberly K. Buhman, and Julie G. Donaldson for their valuable insight. We acknowledge assistance of the Washington University Adipocyte Biology and Molecular Nutrition Core (P30DK056341) and that of the Digestive Disease Research Core (P30DK052574). This work is supported in whole or in part by National Institutes of Health Grants R01 DK088206 (NEW) and DK DK060022 (NAA).

## References

[1] E. Kirk, D.N. Reeds, B.N. Finck, M.S. Mayurranjan, S. Klein, Dietary fat and carbohydrates differentially alter insulin sensitivity during caloric restriction, Gastroenterology, 136 (2009) 1552–1560.

[2] E. Fabbrini, C.T. Luecking, L. Love-Gregory, A.L. Okunade, M. Yoshino, G. Fraterrigo, B.W. Patterson, S. Klein, Physiological Mechanisms of Weight-gain Induced Steatosis in People with Obesity, Gastroenterology, 150 (2016) 79–81.e72.

[3] E. Fabbrini, J. Yoshino, M. Yoshino, F. Magkos, C. Tiemann Luecking, D. Samovski, G. Fraterrigo, A.L. Okunade, B.W. Patterson, S. Klein, Metabolically normal obese people are protected from adverse effects following weight gain, The Journal of Clinical Investigation, 125 (2015) 787–795.

[4] M.Y. Pepino, O. Kuda, D. Samovski, N.A. Abumrad, Structure-Function of CD36 and Importance of Fatty Acid Signal Transduction in Fat Metabolism, Annual review of nutrition, 34 (2014) 281–303.

[5] J.S. Bonifacino, J. Lippincott-Schwartz, Coat proteins: shaping membrane transport, Nature Reviews Molecular Cell Biology, 4 (2003) 409.

[6] S. Mayor, R.G. Parton, J.G. Donaldson, Clathrin-Independent Pathways of Endocytosis, Cold Spring Harbor Perspectives in Biology, 6 (2014) a016758.

[7] M.C. Derby, P.A. Gleeson, New Insights into Membrane Trafficking and Protein Sorting, International Review of Cytology, Academic Press, Place Published, 2007, pp. 47–116.

[8] D.L. Brasaemle, N.E. Wolins, Packaging of fat: an evolving model of lipid droplet assembly and expansion, J Biol Chem, 287 (2012) 2273–2279.

[9] N.E. Wolins, D.L. Brasaemle, P.E. Bickel, A proposed model of fat packaging by exchangeable lipid droplet proteins, FEBS Lett, 580 (2006) 5484–5491.

[10] T. Fujiwara, K. Oda, S. Yokota, A. Takatsuki, Y. Ikehara, Brefeldin A causes disassembly of the Golgi complex and accumulation of secretory proteins in the endoplasmic reticulum, Journal of Biological Chemistry, 263 (1988) 18545–18552.

[11] J. Lippincott-Schwartz, L.C. Yuan, J.S. Bonifacino, R.D. Klausner, Rapid redistribution of Golgi proteins into the ER in cells treated with brefeldin A: Evidence for membrane cycling from Golgi to ER, Cell, 56 (1989) 801–813.

[12] K.G. Soni, G.A. Mardones, R. Sougrat, E. Smirnova, C.L. Jackson, J.S. Bonifacino, Coatomer-dependent protein delivery to lipid droplets, J Cell Sci, 122 (2009) 1834–1841.

[13] D. Hesse, A. Jaschke, B. Chung, A. Schürmann, Trans-Golgi proteins participate in the control of lipid droplet and chylomicron formation, Bioscience Reports, 33 (2013) e00001.

[14] N. Borgese, Getting membrane proteins on and off the shuttle bus between the endoplasmic reticulum and the Golgi complex, Journal of Cell Science, 129 (2016) 1537.

[15] M. Beller, C. Sztalryd, N. Southall, M. Bell, H. Jäckle, D.S. Auld, B. Oliver, COPI Complex Is a Regulator of Lipid Homeostasis, PLoS Biology, 6 (2008) e292.

[16] A.R. Thiam, B. Antonny, J. Wang, J. Delacotte, F. Wilfling, T.C. Walther, R. Beck, J.E. Rothman, F. Pincet, COPI buds 60-nm lipid droplets from reconstituted water–phospholipid– triacylglyceride interfaces, suggesting a tension clamp function, Proceedings of the National Academy of Sciences, 110 (2013) 13244.

[17] E.N. Ellong, K.G. Soni, Q.-T. Bui, R. Sougrat, M.-P. Golinelli-Cohen, C.L. Jackson, Interaction between the Triglyceride Lipase ATGL and the Arf1 Activator GBF1, PLoS ONE, 6 (2011) e21889.

[18] F. Wilfling, A.R. Thiam, M.-J. Olarte, J. Wang, R. Beck, T.J. Gould, E.S. Allgeyer, F. Pincet, J. Bewersdorf, R.V. Farese, T.C. Walther, Arf1/COPI machinery acts directly on lipid droplets and enables their connection to the ER for protein targeting, eLife, 3 (2014) e01607.

[19] S.Y. Ho, L. Delgado, J. Storch, Monoacylglycerol metabolism in human intestinal Caco-2 cells: evidence for metabolic compartmentation and hydrolysis, J Biol Chem, 277 (2002) 1816–1823.

[20] S.Y. Ho, J. Storch, Common mechanisms of monoacylglycerol and fatty acid uptake by human intestinal Caco-2 cells, Am J Physiol Cell Physiol, 281 (2001) C1106–1117.

[21] K.B. Hansen, M.M. Rosenkilde, F.K. Knop, N. Wellner, T.A. Diep, J.F. Rehfeld, U.B. Andersen, J.J. Holst, H.S. Hansen, 2-Oleoyl glycerol is a GPR119 agonist and signals GLP-1 release in humans, J Clin Endocrinol Metab, 96 (2011) E1409–1417.

[22] U. Taschler, F.P.W. Radner, C. Heier, R. Schreiber, M. Schweiger, G. Schoiswohl, K. Preiss-Landl, D. Jaeger, B. Reiter, H.C. Koefeler, J. Wojciechowski, C. Theussl, J.M. Penninger, A. Lass, G. Haemmerle, R. Zechner, R. Zimmermann, Monoglyceride Lipase Deficiency in Mice Impairs Lipolysis and Attenuates Diet-induced Insulin Resistance, The Journal of Biological Chemistry, 286 (2011) 17467–17477.

[23] C.J. Fowler, Monoacylglycerol lipase – a target for drug development?, British Journal of Pharmacology, 166 (2012) 1568–1585.

[24] G. Lyubachevskaya, E. Boyle‐Roden, Kinetics of 2‐monoacylglycerol acyl migration in model chylomicra, Lipids, 35 (2000) 1353–1358.

[25] J.R. Skinner, T.M. Shew, D.M. Schwartz, A. Tzekov, C.M. Lepus, N.A. Abumrad, N.E. Wolins, Diacylglycerol enrichment of endoplasmic reticulum or lipid droplets recruits perilipin 3/TIP47 during lipid storage and mobilization, J Biol Chem, 284 (2009) 30941–30948.

[26] M.C. Gropler, T.E. Harris, A.M. Hall, N.E. Wolins, R.W. Gross, X. Han, Z. Chen, B.N. Finck, Lipin 2 is a liver-enriched phosphatidate phosphohydrolase enzyme that is dynamically regulated by fasting and obesity in mice, J Biol Chem, 284 (2009) 6763–6772.

[27] L.A. Harris, J.R. Skinner, T.M. Shew, T.A. Pietka, N.A. Abumrad, N.E. Wolins, Perilipin 5-Driven Lipid Droplet Accumulation in Skeletal Muscle Stimulates the Expression of Fibroblast Growth Factor 21, Diabetes, 64 (2015) 2757–2768.

[28] N.E. Wolins, B.K. Quaynor, J.R. Skinner, A. Tzekov, C. Park, K. Choi, P.E. Bickel, OP9 mouse stromal cells rapidly differentiate into adipocytes: characterization of a useful new model of adipogenesis, J Lipid Res, 47 (2006) 450–460.

[29] L.A. Harris, J.R. Skinner, N.E. Wolins, Imaging of neutral lipids and neutral lipid associated proteins, Methods Cell Biol, 116 (2013) 213–226.

[30] L.A. Harris, T.M. Shew, J.R. Skinner, N.E. Wolins, A single centrifugation method for isolating fat droplets from cells and tissues, J Lipid Res, 53 1021–1025.

[31] D.M. Schwartz, N.E. Wolins, A simple and rapid method to assay triacylglycerol in cells and tissues, J Lipid Res, 48 (2007) 2514–2520.

[32] G.J. Randolph, N.E. Miller, Lymphatic transport of high-density lipoproteins and chylomicrons, The Journal of Clinical Investigation, 124 (2014) 929–935.

[33] B. Lee, J. Zhu, N.E. Wolins, J.X. Cheng, K.K. Buhman, Differential association of adipophilin and TIP47 proteins with cytoplasmic lipid droplets in mouse enterocytes during dietary fat absorption, Biochim Biophys Acta, 1791 (2009) 1173–1180.

[34] N.E. Wolins, B.K. Quaynor, J.R. Skinner, M.J. Schoenfish, A. Tzekov, P.E. Bickel, S3-12, Adipophilin, and TIP47 package lipid in adipocytes, J Biol Chem, 280 (2005) 19146–19155.

[35] N.E. Wolins, B. Rubin, D.L. Brasaemle, TIP47 associates with lipid droplets, J Biol Chem, 276 (2001) 5101–5108.

[36] C. Xiao, P. Stahel, A.L. Carreiro, K.K. Buhman, G.F. Lewis, Recent Advances in Triacylglycerol Mobilization by the Gut, Trends in Endocrinology & Metabolism, 29 (2018) 151–163.

[37] R. El-Maghrabi, M. Waite, L.L. Rudel, Monoacylglycerol accumulation in low and high density lipoproteins during the hydrolysis of very low density lipoprotein triacylglycerol by lipoprotein lipase, Biochemical and Biophysical Research Communications, 81 (1978) 82–88.

[38] C.J. Fielding, P.E. Fielding, Characteristics of triacylglycerol and partial acylglycerol hydrolysis by human plasma lipoprotein lipase, Biochimica et Biophysica Acta (BBA) - Lipids and Lipid Metabolism, 620 (1980) 440–446.

[39] S.G. Young, R. Zechner, Biochemistry and pathophysiology of intravascular and intracellular lipolysis, Genes & Development, 27 (2013) 459–484.

[40] A.E. Thumser, A.G. Buckland, D.C. Wilton, Monoacylglycerol binding to human serum albumin: evidence that monooleoylglycerol binds at the dansylsarcosine site, J Lipid Res, 39 (1998) 1033–1038.

[41] J. Donaldson, R. Kahn, J. Lippincott-Schwartz, R. Klausner, Binding of ARF and beta-COP to Golgi membranes: possible regulation by a trimeric G protein, Science, 254 (1991) 1197–1199.

[42] M.S. Robinson, T.E. Kreis, Recruitment of coat proteins onto Golgi membranes in intact and permeabilized cells: Effects of brefeldin A and G protein activators, Cell, 69 (1992) 129–138.

[43] E.M. Lynes, T. Simmen, Urban planning of the endoplasmic reticulum (ER): How diverse mechanisms segregate the many functions of the ER, Biochim Biophys Acta.

[44] M. Beller, C. Sztalryd, N. Southall, M. Bell, H. Jackle, D.S. Auld, B. Oliver, COPI complex is a regulator of lipid homeostasis, PLoS Biol, 6 (2008) e292.

[45] D. Hesse, A. Jaschke, B. Chung, A. Schurmann, Trans-Golgi proteins participate in the control of lipid droplet and chylomicron formation, Biosci Rep, 33 (2013) 1–9.

[46] A.R. Thiam, B. Antonny, J. Wang, J. Delacotte, F. Wilfling, T.C. Walther, R. Beck, J.E. Rothman, F. Pincet, COPI buds 60-nm lipid droplets from reconstituted water-phospholipid-triacylglyceride interfaces, suggesting a tension clamp function, Proc Natl Acad Sci U S A, 110 (2013) 13244–13249.

[47] P. Nilsson-Ehle, T. Egelrud, P. Belfrage, T. Olivecrona, B. Borgström, Positional Specificity of Purified Milk Lipoprotein Lipase, Journal of Biological Chemistry, 248 (1973) 6734–6737.

[48] O. Adeyo, C.N. Goulbourne, A. Bensadoun, A.P. Beigneux, L.G. Fong, S.G. Young, GPIHBP1 and the intravascular processing of triglyceride-rich lipoproteins, Journal of internal medicine, 272 (2012) 528–540.

[49] M. Yang, J.T. Nickels, MOGAT2: A New Therapeutic Target for Metabolic Syndrome, Diseases, 3 (2015) 176–192.

[50] O. Adeyo, C.N. Goulbourne, A. Bensadoun, A.P. Beigneux, L.G. Fong, S.G. Young, Glycosylphosphatidylinositol-anchored high-density lipoprotein-binding protein 1 and the intravascular processing of triglyceride-rich lipoproteins, J Intern Med, 272 (2012) 528–540.

[51] R. El Maghrabi, M. Waite, L.L. Rudel, Monoacylglycerol accumulation in low and high density lipoproteins during the hydrolysis of very low density lipoprotein triacylglycerol by lipoprotein lipase, Biochem Biophys Res Commun, 81 (1978) 82–88.

[52] C.J. Fielding, P.E. Fielding, Characteristics of triacylglycerol and partial acylglycerol hydrolysis by human plasma lipoprotein lipase, Biochim Biophys Acta, 620 (1980) 440–446.

[53] P. Nilsson-Ehle, T. Egelrud, P. Belfrage, T. Olivecrona, B. Borgstrom, Positional specificity of purified milk lipoprotein lipase, J Biol Chem, 248 (1973) 6734–6737.

[54] P. de Figueiredo, D. Drecktrah, R.S. Polizotto, N.B. Cole, J. Lippincott-Schwartz, W.J. Brown, Phospholipase A2 antagonists inhibit constitutive retrograde membrane traffic to the endoplasmic reticulum, Traffic, 1 (2000) 504–511.

[55] E. Ikonen, Roles of lipid rafts in membrane transport, Curr Opin Cell Biol, 13 (2001) 470–477.

[56] E. Orso, C. Broccardo, W.E. Kaminski, A. Bottcher, G. Liebisch, W. Drobnik, A. Gotz, O. Chambenoit, W. Diederich, T. Langmann, T. Spruss, M.F. Luciani, G. Rothe, K.J. Lackner, G. Chimini, G. Schmitz, Transport of lipids from golgi to plasma membrane is defective in tangier disease patients and Abc1-deficient mice, Nat Genet, 24 (2000) 192–196.

[57] J.A. Schmidt, W.J. Brown, Lysophosphatidic acid acyltransferase 3 regulates Golgi complex structure and function, J Cell Biol, 186 (2009) 211–218.

[58] A. Simonsen, A.E. Wurmser, S.D. Emr, H. Stenmark, The role of phosphoinositides in membrane transport, Curr Opin Cell Biol, 13 (2001) 485–492.

[59] K. Chambers, W.J. Brown, Characterization of a novel CI-976-sensitive lysophospholipid acyltransferase that is associated with the Golgi complex, Biochem Biophys Res Commun, 313 (2004) 681–686.

[60] O. Adeyo, P.J. Horn, S. Lee, D.D. Binns, A. Chandrahas, K.D. Chapman, J.M. Goodman, The yeast lipin orthologue Pah1p is important for biogenesis of lipid droplets, J Cell Biol, 192 1043–1055.

[61] N. Ariotti, S. Murphy, N.A. Hamilton, L. Wu, K. Green, N.L. Schieber, P. Li, S. Martin, R.G. Parton, Postlipolytic insulin-dependent remodeling of micro lipid droplets in adipocytes, Mol Biol Cell, 23 1826–1837.

[62] V. Litvak, Y.D. Shaul, M. Shulewitz, R. Amarilio, S. Carmon, S. Lev, Targeting of Nir2 to Lipid Droplets Is Regulated by a Specific Threonine Residue within Its PI-Transfer Domain, Current Biology, 12 (2002) 1513–1518.

[63] J.A. Schmidt, W.J. Brown, Lysophosphatidic acid acyltransferase 3 regulates Golgi complex structure and function, The Journal of Cell Biology, 186 (2009) 211.

[64] A. Simonsen, A.E. Wurmser, S.D. Emr, H. Stenmark, The role of phosphoinositides in membrane transport, Current Opinion in Cell Biology, 13 (2001) 485–492.

[65] J. Scheel, R. Pepperkok, M. Lowe, G. Griffiths, T.E. Kreis, Dissociation of Coatomer from Membranes Is Required for Brefeldin A–induced Transfer of Golgi Enzymes to the Endoplasmic Reticulum, The Journal of Cell Biology, 137 (1997) 319–333.

[66] M. Langhans, D.G. Robinson, 1-Butanol targets the Golgi apparatus in tobacco BY-2 cells, but in a different way to Brefeldin A, Journal of Experimental Botany, 58 (2007) 3439–3447.

[67] C. Dascher, W.E. Balch, Dominant inhibitory mutants of ARF1 block endoplasmic reticulum to Golgi transport and trigger disassembly of the Golgi apparatus, Journal of Biological Chemistry, 269 (1994) 1437–1448.

[68] O. Stein, Y. Dabach, G. Hollander, M. Ben-Nairn, Y. Stein, Dissimilar effects of Brefeldin A on cholesteryl ester and triacylglycerol metabolism in CaCo2 and HepG2 cells as compared to peritoneal macrophages, Biochimica et Biophysica Acta (BBA) - Lipids and Lipid Metabolism, 1125 (1992) 28–34.

[69] Y. Ohsaki, J. Cheng, M. Suzuki, A. Fujita, T. Fujimoto, Lipid droplets are arrested in the ER membrane by tight binding of lipidated apolipoprotein B-100, Journal of Cell Science, 121 (2008) 2415.

